# ProteoSushi: a software tool to biologically annotate and quantify modification-specific, peptide-centric proteomics datasets

**DOI:** 10.1101/2020.11.24.395921

**Authors:** Robert W. Seymour, Sjoerd van der Post, Arshag D. Mooradian, Jason M. Held

## Abstract

Large-scale proteomic profiling of protein post-translational modifications has provided important insights into the regulation of cell signaling and disease. These modification-specific proteomics workflows nearly universally enrich modified peptides prior to mass spectrometry analysis, but protein-centric proteomic software tools have many limitations evaluating and interpreting these peptide-centric datasets. We therefore developed ProteoSushi, a software tool tailored to the analysis of each modified site in peptide-centric proteomic datasets that is compatible with any post-translational modification or chemical label. ProteoSushi uses a unique approach to assign identified peptides to shared proteins and genes, minimizing redundancy by prioritizing shared assignments based on UniProt annotation score and optional user-supplied protein/gene lists. ProteoSushi simplifies quantitation by summing or averaging intensities, merging overlapping peptide charge states, missed cleavages, peptide spectral matches, and variable modifications into a single value for each modified site. ProteoSushi annotates each PTM site with the most up-to-date biological information available from UniProt, such as functional roles or known modifications, the protein domain in which the site resides, the protein’s subcellular location and function and more. ProteoSushi has a graphical user interface for ease of use. ProteoSushi’s flexibility and combination of features streamlines peptide-centric data processing and knowledge mining of large modification-specific proteomics datasets.

## Introduction

Protein post-translational modifications (PTMs) expand the proteome’s function by regulating protein structure, activity, localization, and interactions. Approximately 5% of human genes encode PTM modifying enzymes, and of the several million proteoforms found within a cell, the vast majority of proteome diversity stems from PTMs^1^. There are hundreds of known PTMs, with some PTMs predominantly regulated by enzyme activity and others more often the result of chemical modification^2,3^.

Large-scale, unbiased identification and quantitation of modified residues using proteomics has provided unparalleled insight into the regulation of PTM networks, typically quantifying many thousands of modified sites in parallel. In a typical modification-focused proteomic experiment, the PTM of interest is enriched via antibodies or affinity tags^2,4,5^. Transient modifications can be also be chemically labeled for chemoproteomics, potentially utilizing click chemistry to conjugate on a handle for enrichment such as biotin^6,7^. Thus, a key feature of modification-specific proteomics datasets is a highly enriched set of peptides modified by the target modification(s). There is typically a single target PTM or chemical label of interest per peptide, or several in the case of differential alkylation of cysteines or phosphorylation of serine, threonine, and tyrosine^8,9^.

The data processing goals of modification-specific datasets coupled with their peptidecentric nature dictate several unique analytical challenges compared to protein-centric datasets. First, typical database search algorithms leverage the many peptides identified in a sample to infer which related protein isoforms or protein groups are present, and then assign peptides to these protein groups parsimoniously. However, protein coverage in modification-specific proteomic datasets is sparse, as only modified peptides are analyzed. Since each modified peptide identified is a unique entity, it must be treated as such for analysis. Assigning modified peptides using typical protein-based typical grouping algorithms is inconsistent and varies depending on the other peptides identified in a sample. Grouping modified peptides at the protein level may inappropriately minimize the actual level of ambiguity in protein assignment, especially since the amino acids proximal to regulatory PTMs are often highly conserved^10–12^. Second, database search tools don’t provide downstream annotation of the modified sites identified. Third, multiple peptide modifications detected on the modified peptides due to sample processing, such as deamidation or methionine oxidation, or missed cleavages, are of limited biological interest. Therefore, the typical goal of modification-specific analysis is to collapse these additional peptides to a single PTM site and quantitative value for simplification, a feature not typical in protein-centric tools that focus on grouping proteins rather than PTM sites.

An array of software tools exists for peptide- and PTM-centric analysis including databases^13–18^, PTM site assignment^19–22^, and visualization^23,24^. However, no software tool is able to quantify and annotate any site and type of modified residue(s) in proteomic datasets. Software tools focused on downstream processing of proteomic data include Perseus, an integrated tool for statistics and comparisons across samples^25^. However, Perseus is in essence protein-centric and has limited capacity to annotate all but the most common PTM sites with up-to-date information. PTMsigDB is another PTM-focused annotation tool to identify biological signatures based on site-specific, rather than gene-level data, but it is limited to phosphorylation and is also designed to act downstream of many key peptide-centric processing decisions and steps^26^. Skyline^27^, a software tool for label free quantitation and analysis of data-independent acquisition (DIA) data^28^, is peptide-focused, though the resulting output requires reassignment of modified peptides to shared proteins and genes, merging of modified peptides and charge states, and biological annotation of the modified sites^29^. Therefore, there is an unmet need for a flexible and easy to use pipeline to process, characterize, and interrogate modification-specific proteomic data downstream of both database search and quantitative results.

To meet this need we developed ProteoSushi, a purpose-built software framework tailored to the peptide centric quantification, protein assignment and biological annotation of modification-specific proteomics data (**Fig. 1**). ProteoSushi is highly flexible and can be applied to any PTM or chemical mass addition of interest, and can be used for any type of quantitation, including isotopic- and isobaric-labeling and data-independent acquisition (DIA) datasets. ProteoSushi integrates quantitative information from all forms of a modified peptide covering the same site including charge states, missed cleavages, and other variable modifications such as methionine oxidation and asparagine deamidation found in the peptide sequence, to a single value, greatly simplifying statistics and interpretation. ProteoSushi assigns each peptide to all shared proteins and genes based on the peptide spectral match search database output or peptide list, prioritizing those with higher UniProt annotation scores for the most comprehensive annotation to balance the inherent ambiguity of assignment to proteins and genes that share the peptide sequence. In addition, ProteoSushi can prioritize user-defined lists of proteins for situations such as enrichment of a specific organelle where a subset of homologous proteins may be expected to be present. For quantitation, ProteoSushi imports the quantitative results from search engine outputs or DIA-MS, and can either sum or average the intensities. Lastly, ProteoSushi annotates each modified peptide based on the most up-to-date UniProt knowledgebase^30^ retrieved in real time via a web API. This includes residue-specific annotations such as known PTMs, metal binding, or mutagenesis; protein domain-level annotations such as secondary structure and the protein domain the modification resides in; and protein-level annotation such as function and subcellular location.

**Figure 1.**
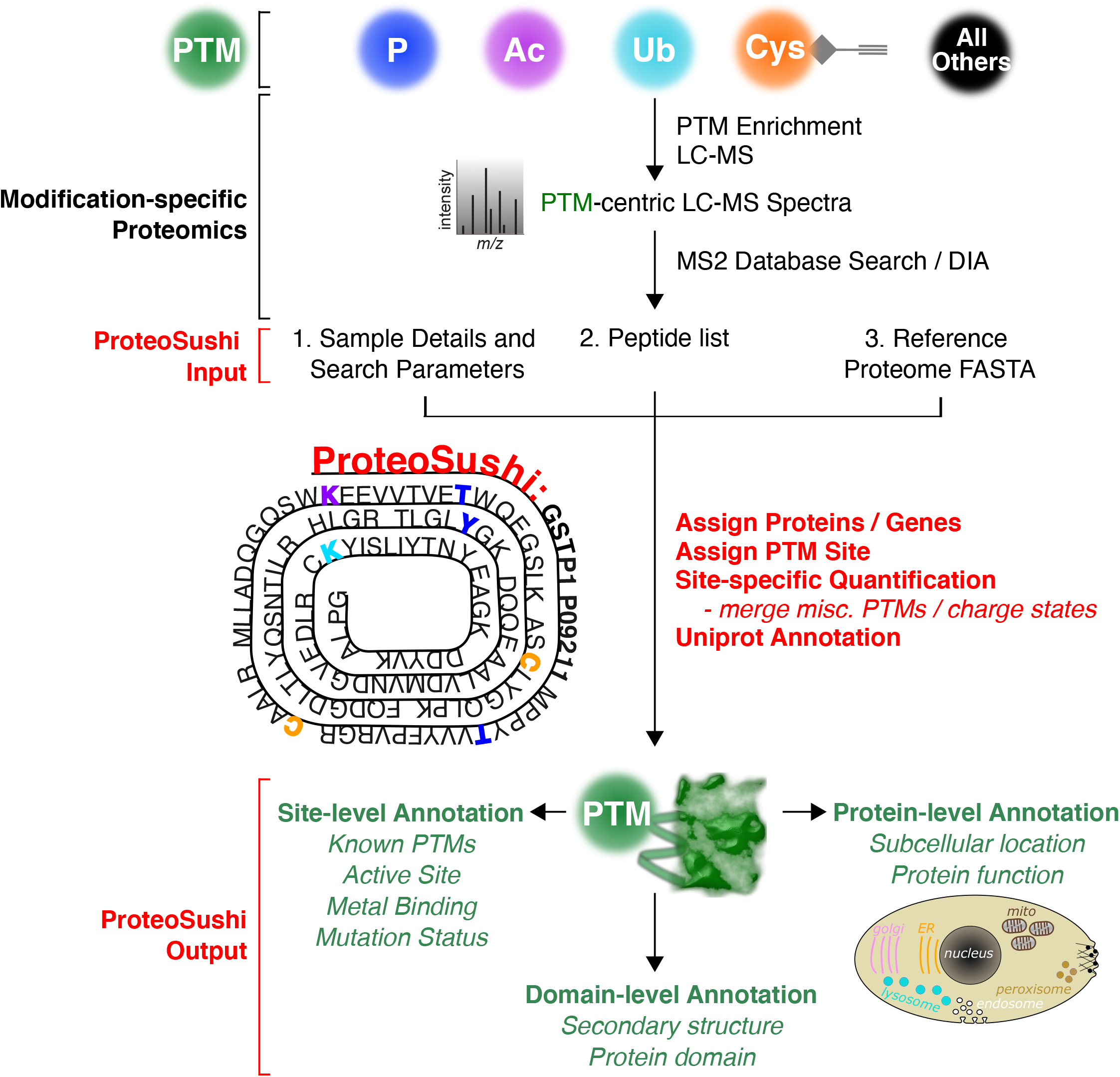
ProteoSushi overview. ProteoSushi processes modification-specific proteomics datasets typically enriched for peptides with a single, or a small number, of PTMs or chemical modifications. This includes phosphorylation (P), acetylation (Ac), ubiquitylation (Ub), as well as chemical trapping of cysteine modifications. ProteoSushi requires several input files, including a list of peptides with modifications, and a proteome FASTA file. ProteoSushi then assigns PTMs to shared proteins and genes, reducing complexity by combining forms of the peptides sharing the same PTMs with additional chemical modifications, cleavage sites, or charge states. The UniProt SPARQL endpoint is used to provide site-, domain-, and protein-specific biological annotations.

ProteoSushi is freely available at http://github.com/HeldLab/ProteoSushi and utilizes a graphical user interface (GUI) for ease of use (**Fig. 2**). A computer running ProteoSushi needs at least 8 gigabytes of RAM (12 gigabytes on a Windows-based system), Python, and a stable internet connection to retrieve UniProt information. A detailed online tutorial including step-by-step instructions is available to guide the user through the installation and use on the ProteoSushi github (https://github.com/HeldLab/ProteoSushi). Example files are included as well (https://tinyurl.com/y5gtk8y9). A flow chart detailing each assignment step and quantitation integration of modified peptides is included as **Supp. Fig. S1**.

**Figure 2.**
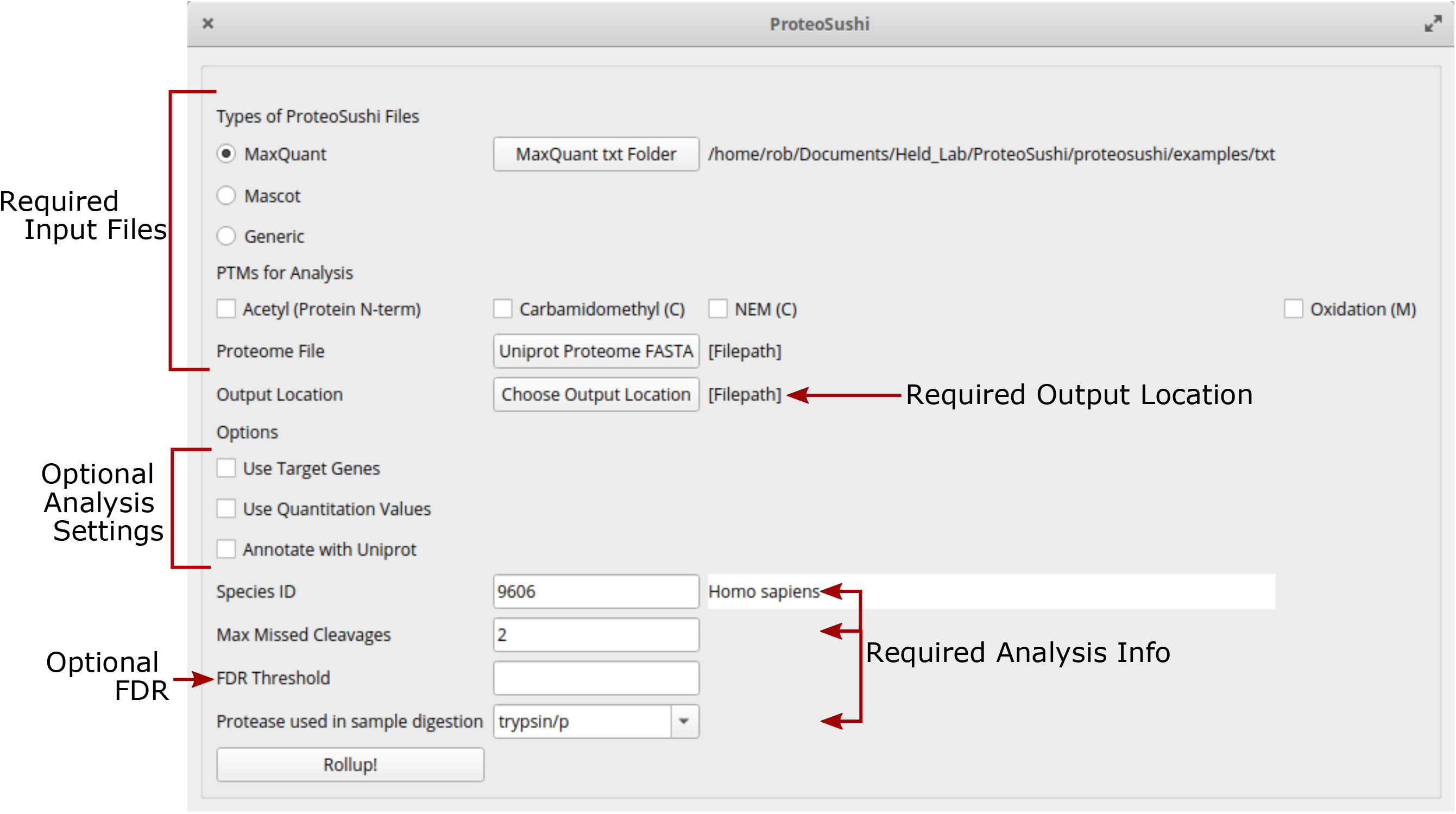
ProteoSushi graphical user interface (GUI). The GUI includes options to analyze generic peptide lists, or directly process the output files from MaxQuant or Mascot. A prompt enables loading of the proteome FASTA file and flexible options for the user to tailor processing based on species, quantitation, peptide false discovery rate filtering and other options.

## Results and Discussion

### Identification: Parsing the modifications of interest and assigning modified peptides to proteins, genes and sites

ProteoSushi requires two input files: 1) a FASTA file, typically the same file used for the database search such as a UniProt reference proteome, and 2) a list of peptides, both with and without modifications, such as the modified peptide sequences detected by a database search (**Fig. 2**). Alternatively, the peptide lists, from Skyline or from a supplementary table for example, can be used as input for annotation and quantification. Quantative values can be optionally included. The unrestricted nature of ProteoSushi inputs gives it the flexibility to be used for any peptide-centric dataset including any species and any database search tool. For convenience, ProteoSushi is designed to directly accept input from the output files from MaxQuant^31^ and Mascot^32^, two widely used database search engines, but also accepts any CSV file with columns for modified and unmodified peptides. Specific information about this format is provided on the GitHub page (https://github.com/HeldLab/ProteoSushi).

ProteoSushi can process any mass shift(s) and modified residue(s), and thus can be applied to any PTM or chemical modification of interest including cases where several modifications are of interest, such as phosphorylation which can occur on serine, threonine, or tyrosine^9^. ProteoSushi can also be used for chemoproteomic results, as in the case of differential alkylation of cysteines, where reduced and oxidized cysteines are labeled with distinct chemical reagents^33^. Each modification present in the peptide list is retrieved by ProteoSushi and presented as an option in the GUI to select as a modification target. Additional modifications that are not the focus, such as chemical artifacts or routine cysteine alkylation, are merged together for quantitation along with missed cleaved and multiply charged forms of the peptide.

ProteoSushi assigns each peptide to all proteins with a matching sequence in the user-provided FASTA proteome using the amino acid sequence of each modified peptide. Leucines and isoleucines, which are isobaric and can’t be distinguished through mass spectrometry, are considered equivalent. Proteins in UniProt reference proteome are not equally likely studied and characterized resulting in a wide range of annotation quality and depth. Therefore, ProteoSushi compares the UniProt annotation scores of all matched proteins and annotates the peptide with the available information of those with the highest score (**Supp. Fig. S1**). This approach minimizes over-annotation and prioritizes assignment to better characterized proteins. This is especially useful for species that are not annotated well. The primary gene name assigned to the UniProt ID is used to assign the gene, and the modified residue(s) of the peptide are assigned to the amino acid number in the UniProt amino acid sequence.

ProteoSushi provides an option for a user-input list of gene names to prioritize identification to a specific subset of candidate proteins. Potential applications of this include organelle or protein complex enrichment prior to PTM enrichment, such as mitochondrial fractionation followed by enrichment for lysine acetylation^4^. In this case, the user can input a list of mitochondrial genes and ProteoSushi will prioritize assignment of peptides to these genes instead of cytoplasmic proteins that may have sequence homology. The ProteoSushi output indicates that the assigned proteins are in the target list to simplify follow-up analysis.

To compare how proteins and genes are assigned by ProteoSushi versus typical protein-centric database search software, the data output from MaxQuant were compared before and after ProteoSushi processing. A publicly available proteomic dataset that enriched and quantified the reversible oxidation of ~4,000 cysteines^29^ was used for evaluation. Focusing on the carbamidomethylated peptide YDGYYTSC(ca)PLVTGYYNR, and its missed cleaved form KYDGYYTSC(ca)PLVTGYYNR, the MaxQuant output has more than twelve rows of information for different peptide spectral matches (PSMs), including different charge states (**Fig. 3**). Typical database search results also include additional peptide forms with chemical modifications. The peptide is assigned to two proteins with UniProt IDs H3BNP9 and Q9Y6N5, with UniProt annotation scores of 1 and 5, respectively, indicating that the level of annotation of Q9Y6N5 is better. However, since MaxQuant uses the identified proteins to prioritize protein assignment, the poorly annotated H3BNP9 is indicated as the leading protein. In contrast, ProteoSushi lists Q9Y6N5 as the assigned protein based on its higher UniProt annotation score. ProteoSushi also provides the gene name, site assignment of the modified residue and minimizes redundancy by collapsing missed cleaved peptides, multiple charge states, and chemical modifications down to a single row (**Fig. 3**). In addition, ProteoSushi biologically annotates the modified site and protein. Notably, C379 is only annotated in a disulfide bond in Q9Y6N5 as opposed to H3BNP9, highlighting the benefits of annotation score prioritization.

**Figure 3.**
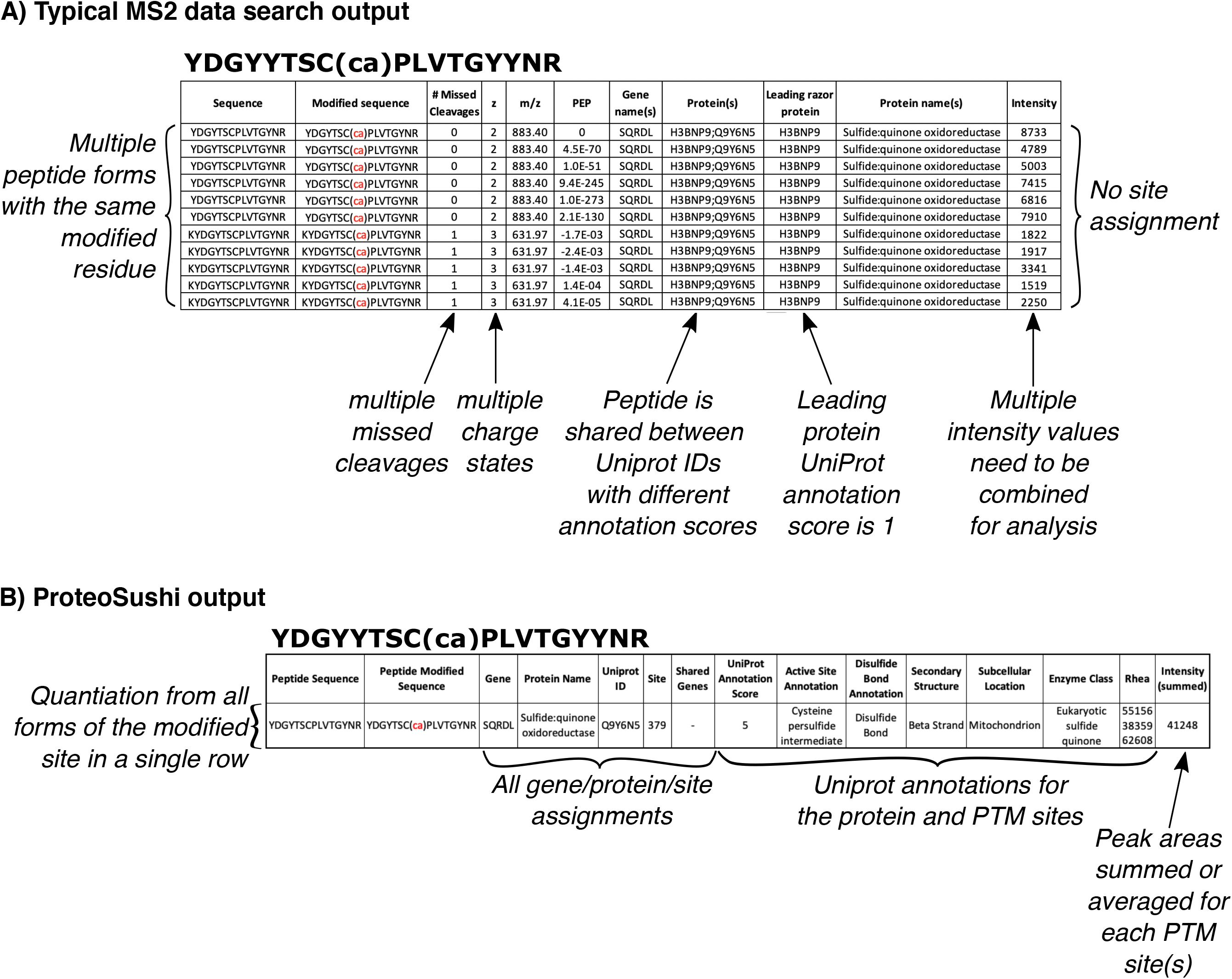
Quantitative proteomics data processing with ProteoSushi. **A)** A typical search result from a protein-centric database search engine such as MaxQuant for the carbamidomethylated (ca) peptide YDGYYTSC(ca)PLVTGYYNR, and its missed cleaved form KYDGYYTSC(ca)PLVTGYYNR. Each peptide spectral match is listed in a new row. **B)** ProteoSushi output that collapses peptide forms with different charge states and missed cleavages, biological annotation from UniProt, and intensity values from each PSM combined into a single value.

### Site-specific data quantitation for each modified residue

ProteoSushi simplifies the quantitative results by integrating results from all peptide modified sequences that have the same PTM site(s) such as multiple charges, missed cleaved peptides, and additional variable modifications such as methionine oxidation or deamidation (**Fig. 3**). The user has an option to sum the areas, which would be appropriate for DIA results, isotopic labelling and label free quantification, or averaging the results, which is a useful option for the reporter ion ratios reported by isobaric tagging studies. Quantitative results are reported for each identified PTM site (**Fig. 3**).

### Biological annotation of PTM sites: Mining knowledge at the protein-, domain-, and sitespecific level

Database search engines do not provide annotation of the identified proteins, and downstream proteomic tools that are protein-centric do not provide site-specific biological annotations (**Fig. 3**). Up-to-date annotation allows researchers to make the best use of current knowledge for scientific discoveries. For example, it was demonstrated that in the majority of the biological pathway analysis studies the full biological impact was missed due to outdated gene annotation used by the analysis tools^34^. ProteoSushi facilitates knowledge mining of modification-specific datasets by using UniProt’s SPARQL endpoint to provide residue- and protein-specific information for each identified site (**Fig. 4**). This approach enables ProteoSushi to provide up to date annotations on the fly for any of the nearly 200,000,000 entries in UniProt expressed in more than 1,000,000 species. Site-specific annotated features include whether the site is known to be a modified active site, binds known ligands, has functional consequences when mutated and others (**Fig. 4A**). Domain-level annotations include the domain type, secondary structure and topology that the modified site resides in. Protein-level annotations include subcellular location and function. UniProt includes known annotations for many different PTMs as well (**Fig. 4B**). Many these are rapidly changing, such as succinyl lysine annotation in human UniProt entries (**Fig. 4C**), despite no increase in total human protein entries in UniProt over time (**Fig. 4D**), underscoring the importance of utilizing updated annotations. Similar increases in UniProt annotations over time are observed for many residue and protein-level annotations (**Fig. 4E**), including protein secondary structure (**Fig. 4F**).

**Figure 4.**
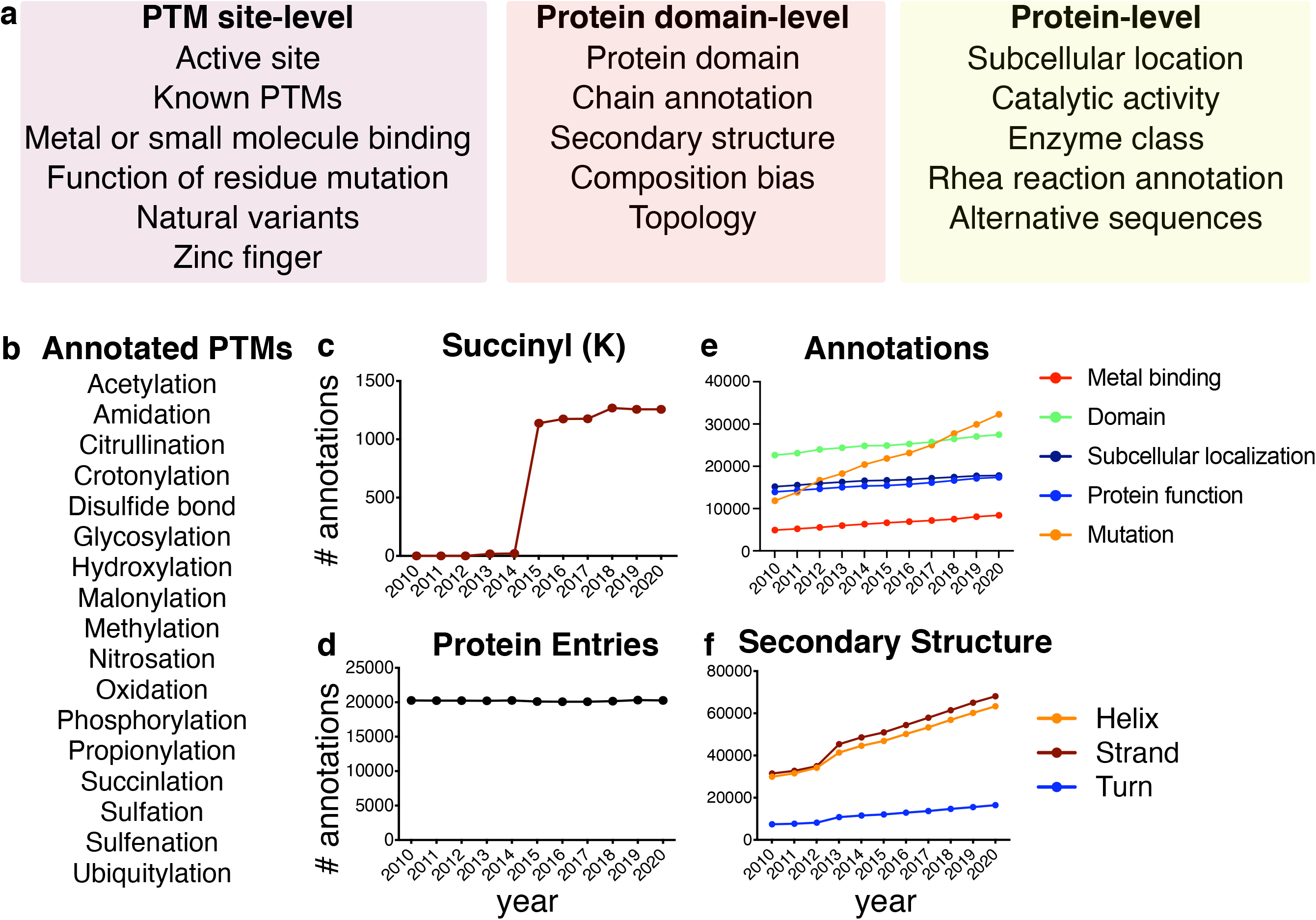
ProteoSushi retrieves UniProt annotations for PTM-sites and modified proteins. **A)** ProteoSushi retrieves UniProt Annotations on the fly via the SPARQL endpoint for the PTM site-, protein domain-, and protein-level biological annotations indicated. **B)** A subset of the PTMs annotated in UniProt. **C)** Number of annotated human UniProt (Swiss-Prot) entries for succinyl lysine, **D)** total human protein entries in UniProt (Swiss-Prot), **E)** miscellaneous UniProt (Swiss-Prot) annotations for proteins and sites, including annotation of mutagenesis, and **F)** protein secondary structure.

## Conclusions

We report ProteoSushi, an open source, platform independent software tool that assigns, quantifies, and annotates peptide-centric proteomics data in a single tool. ProteoSushi fills the void between database search engines and discovery. It is focused on user flexibility, accepting generic peptide lists as input and thus is compatible with any database search engine. It can be applied to all peptide modifications and species, and can process data from all common quantitative mass spectrometry platforms. ProteoSushi complements the many PTM databases and PTM-focused tools available for functional prediction^35^ and text mining^36,37^. ProteSushi can be implemented in any lab and operating system, and has a GUI for ease of use. By facilitating knowledge extraction from modification-specific datasets, ProteoSushi enables better understanding of the function of PTMs and the regulation of signal transduction networks.

## Acknowledgements

We acknowledge funding from R01 CA200893 (J.H). SVDP is supported by the SSMF foundation (P17-0060) We acknowledge Tristaan Haddad for evaluating ProteoSushi output files.

## Supporting Info

**Supplemental Figure S1. Rules for peptide assignment and merging multiple peptide forms.**

Flow chart detailing how ProteoSushi assigns peptides to shared proteins and genes as well as combines multiple peptide forms.

## Abbreviations

PTM: post-translational modification
GUI: graphical user interface
DIA: data-independent acquisition
API: application programming interface
FASTA: fast all
PSM: peptide spectral match
SPARQL: SPARQL Protocol and RDF Query Language
P: phosphorylation
Ac: acetylation
Ub: ubiquitylation
FDR: false discovery rate

