## Supplementary figures and images for "ProteoSushi: a software tool to biologically annotate and quantify modification-specific, peptide-centric proteomics datasets"

### Supplemental Figure 1

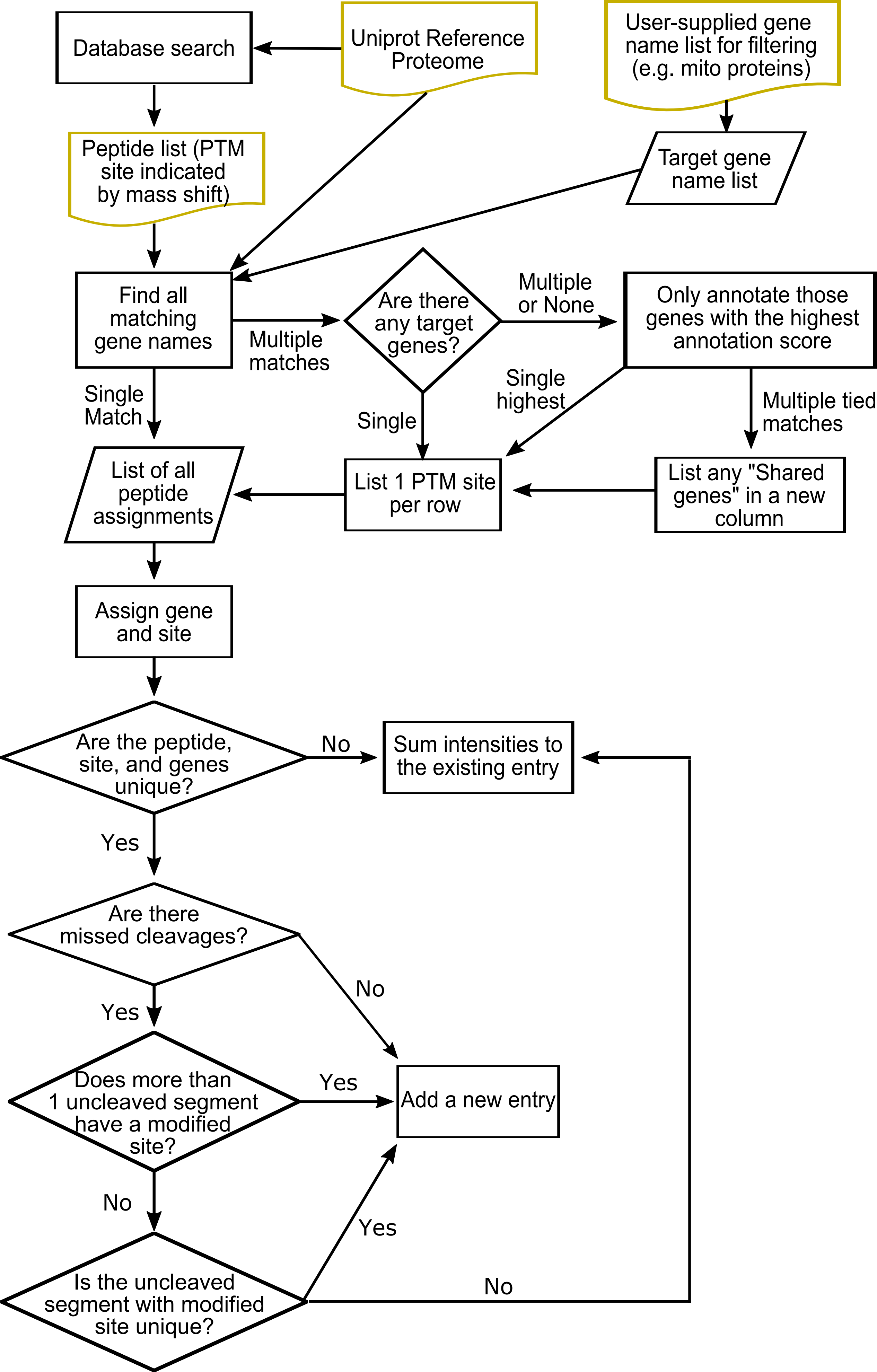
